# Molecular and Structural Reprogramming of Gastric Cancer Revealed by Systems-Level Transcriptomic Analysis

**DOI:** 10.64898/2026.02.18.706548

**Authors:** Negar Mottaghi-Dastjerdi, Mohammad Soltany-Rezaee-Rad

**Affiliations:** Department of Pharmacognosy and Pharmaceutical Biotechnology, School of Pharmacy, Iran University of Medical Sciences, Tehran, Iran; Behestan Innovation Factory, Behestan Darou, Tehran, Iran

**Author notes:** Corresponding Author: Dr. Negar Mottaghi-Dastjerdi, Department of Pharmacognosy and Pharmaceutical Biotechnology, School of Pharmacy, Iran University of Medical Sciences, Tehran, Iran.

**Keywords:** Biomarker discovery, Gastric cancer, Gene network analysis, HOX genes, Systems pathology, Transcriptomic profiling

## Abstract

Gastric cancer (GC) is a major cause of cancer mortality and remains difficult to diagnose early and treat effectively. Although transcriptomic profiling has defined extensive molecular heterogeneity, many studies are not anchored to clinicopathological variables or interpreted in the context of tissue-level pathobiology. We applied an integrative transcriptomic and network-based framework to identify molecular signatures that reflect structural and biochemical reprogramming of gastric tissue during malignant transformation. RNA-seq expression profiles and clinical annotations were analyzed using non-parametric differential expression filtering, functional enrichment, and protein–protein interaction network modeling. Expression patterns were further evaluated across clinicopathological strata (stage, grade, nodal status, and metastasis) and by network-informed clustering. Across clinical strata, GC showed a consistent signature of developmental reactivation and loss of gastric epithelial identity. Developmental regulators, most prominently HOX-cluster genes and the histone variant HIST1H3J, were upregulated, consistent with epigenetic/chromatin reprogramming. In parallel, gastric differentiation and secretory lineage markers (ATP4A, KCNE2, PTF1A, VSTM2A) were persistently downregulated, reflecting suppression of parietal/ductal programs and altered metabolic/secretory function. ADIPOQ demonstrated stage-dependent repression and was associated with poorer survival, supporting a context-dependent prognostic role. Enrichment and network analyses also highlighted FGFR-centered signaling as a dominant oncogenic axis linked to proliferation and invasion. This study identifies a molecular pathology framework in which GC progression involves coordinated chromatin-level developmental reprogramming and sustained loss of gastric differentiation programs, accompanied by FGFR-driven oncogenic signaling and stage-dependent metabolic disruption. These signatures provide candidate diagnostic and prognostic biomarkers and support prioritization of therapeutically actionable pathways in GC.

## 1. Introduction

Gastric cancer (GC) remains as one of the main global health burden and ranks among the leading causes of cancer-related mortality, especially in East Asian populations where late-stage diagnoses and therapeutic resistance are prevalent [1, 2]. Recent data indicate that GC remains highly prevalent across East Asia, with persistent high mortality and only modest improvements in survival over recent years [1, 3, 4]. Despite advances in surgical and chemotherapeutic intervention, the 5-year overall survival of GC remains below 30-35% throughout much of the world, particularly with diagnoses at advanced stages [1, 5]. In the last decade, large-scale transcriptomic profiling analysis, utilizing large datasets including those based on TCGA data and integrated bulk/single-cell datasets, has illuminated extensive molecular heterogeneity and complexity in GC, uncovering novel subtypes, signaling networks, and gene expression patterns [6, 7]. However, it is yet difficult to translate those findings into clinically valuable biomarkers and therapeutic targets.

GC pathobiology involves coordinated transcriptional dysregulation, immune evasion, metabolic rewiring, and dedifferentiation. During tumorigenesis, gastric epithelial identity is progressively lost as embryonic/developmental pathways (e.g., Wnt, Notch, FGFR) are reactivated, producing transcriptomic signatures linked to early malignancy and clinical outcomes. Systems-biology and gene-network analyses help decode these complex patterns and prioritize central regulatory genes with translational relevance [8, 9].

Despite extensive transcriptomic profiling, many studies still report DEGs without network context or stratification by key clinicopathological variables (stage, grade, nodal status). As a result, systems-level insight into developmental reactivation (e.g., HOX clusters) and loss of epithelial lineage programs remains limited. While integrative network modeling has proven useful for identifying drivers and resistance mechanisms across GI and hepatic cancers [10–16], unified pipelines that link phenotype correlations with multidimensional network clustering are still uncommon, leaving a gap at the intersection of transcriptomics, network centrality, and clinical relevance for biomarker discovery.

To address these gaps, we performed an integrated transcriptomic and systems-level analysis of TCGA-STAD RNA-seq data, combining non-parametric DEG filtering with dual PPI network construction, functional enrichment, and clinicopathological stratification (stage, grade, nodal status). This framework aimed to identify stable diagnostic and prognostic biomarkers, characterize developmental and immune transcriptional programs, and prioritize therapeutic candidates, particularly within HOX genes, gastric lineage markers, and core regulatory transcription factors—building on our prior work in gastric and other gastrointestinal cancers [17, 18].

## 2. Materials and methods

### 2.1. Computational environment and tools

All preprocessings of RNA-seq data, normalization, filtering, and differential expression analysis were carried out in Python (v3.10) via Jupyter Notebook. Main involved libraries were pandas for data manipulation, numpy for mathematical calculations, and matplotlib and seaborn for data plotting purposes.

### 2.2. Filtering low-expression genes and log2 normalization

Raw counts of 448 TCGA gastric cancer samples were removed for low-expression genes (≤1 count in >95% of the samples), and 56,678 genes expressed in ≥5 samples were left. Log₂(x + 1) transformation was then applied for stabilization of variance and for enhancing comparability.

### 2.3. Clinical metadata processing

Clinical data for 511 TCGA-STAD samples were retrieved from GDC, and key variables (e.g., ID, gender, stage, survival) were extracted. Data were cleaned and standardized using Python (v3.10) and pandas.

### 2.4. Differential expression analysis (DEA)

RNA-seq data from 448 samples (412 tumors and 36 normal tissues) were analyzed using the Mann–Whitney U test. Differentially expressed genes (DEGs) were defined as those with |log2FC| ≥ 1 and FDR < 0.05 (Benjamini–Hochberg). Genes with an average expression ≥ 1 in both groups were retained, and the top 100 upregulated and top 100 downregulated DEGs were selected for downstream analyses. Ensembl IDs were converted to gene symbols using MyGene.info. All analyses were performed in Python (v3.10) using the pandas, scipy, statsmodels, and mygene packages. The complete DEG list is provided in **Supplementary File 1**.

### 2.5. Functional enrichment analysis

Upregulated (n=85) and downregulated (n=94) DEGs were analyzed using ENRICHR to identify enriched Gene Ontology (GO: Biological Process (BP), Molecular Function (MF), and Cellular Component (CC)) terms and Reactome 2024 pathways. Only results with FDR < 0.05 were considered significant. Enrichment results were visualized as bar plots based on −log₁₀(FDR) for comparative interpretation.

### 2.6. Protein–protein interaction (PPI) network analysis

Two PPI networks were constructed using STRING in Cytoscape v3.10: one from ∼6,500 DEGs (split into four subsets) and another from the top 100 up- and downregulated genes. Networks were generated with a confidence cutoff of 0.15. Hub genes were identified using cytoHubba via MCC, DMNC, MNC, and Degree methods; genes ranked top-5 across multiple metrics were deemed high-confidence hubs.

### 2.7. Cluster analysis of the network

To identify functional modules within the 6,500-gene PPI network, we applied the IPCA algorithm in CytoCluster (Cytoscape v2.1.0) [19] with TinThreshold = 0.9, ComplexSize ≥ 10, and PathLength = 2. This yielded 28 clusters. The top 4 clusters (ranked by density) were analyzed for pathway enrichment using Reactome via Enrichr, focusing on the top 20 significant pathways per cluster.

### 2.8. UALCAN-based clinicopathological expression and survival analysis

Expression of the 13 hub genes was assessed in TCGA-STAD using the UALCAN portal (https://ualcan.path.uab.edu). TPM values were retrieved for cancer stage, tumor grade, nodal metastasis status, and survival modules, with normal gastric tissue as the reference. UALCAN applies Welch’s unpaired t-tests for subgroup comparisons, and Kaplan–Meier curves were used to evaluate prognostic associations (P < 0.05).

## 3. Results

### 3.1. Identification of DEGs in GC

RNA-seq data of 448 gastric samples (412 tumor, 36 normal) was compared with the Mann–Whitney U test to identify the DEGs. The genes with a threshold of |log2FC| ≥ 1 and FDR < 0.05 were designated as significant, and there were 9,896 DEGs (9,592 upregulated, 304 downregulated). To improve reliability, we retained only genes with average expression ≥ 1 in both groups. From the filtered list, the top 100 upregulated and top 100 downregulated DEGs (by absolute log2FC) were taken for downstream analysis. Gene IDs were converted to official symbols using the MyGene.info API.

### 3.2. Network reconstruction and analysis

To distinguish therapeutic targets from diagnostic/prognostic biomarkers, we constructed two PPI networks from TCGA-STAD DEGs. The genome-scale network (∼6,500 DEGs) mapped global GC interactions and, using cytoHubba (four topology algorithms), identified highly central hubs, including histone variants (H3C12, H3-4), transcription factors (PAX2, HOXD11/13, HOXC12), signaling molecules (PRKACG, FGF8), and immune mediators (IFNG, IFNL2), as candidate drug targets because perturbing them could disrupt broad tumor-promoting circuitry (**Table 1**).

**Table 1.**
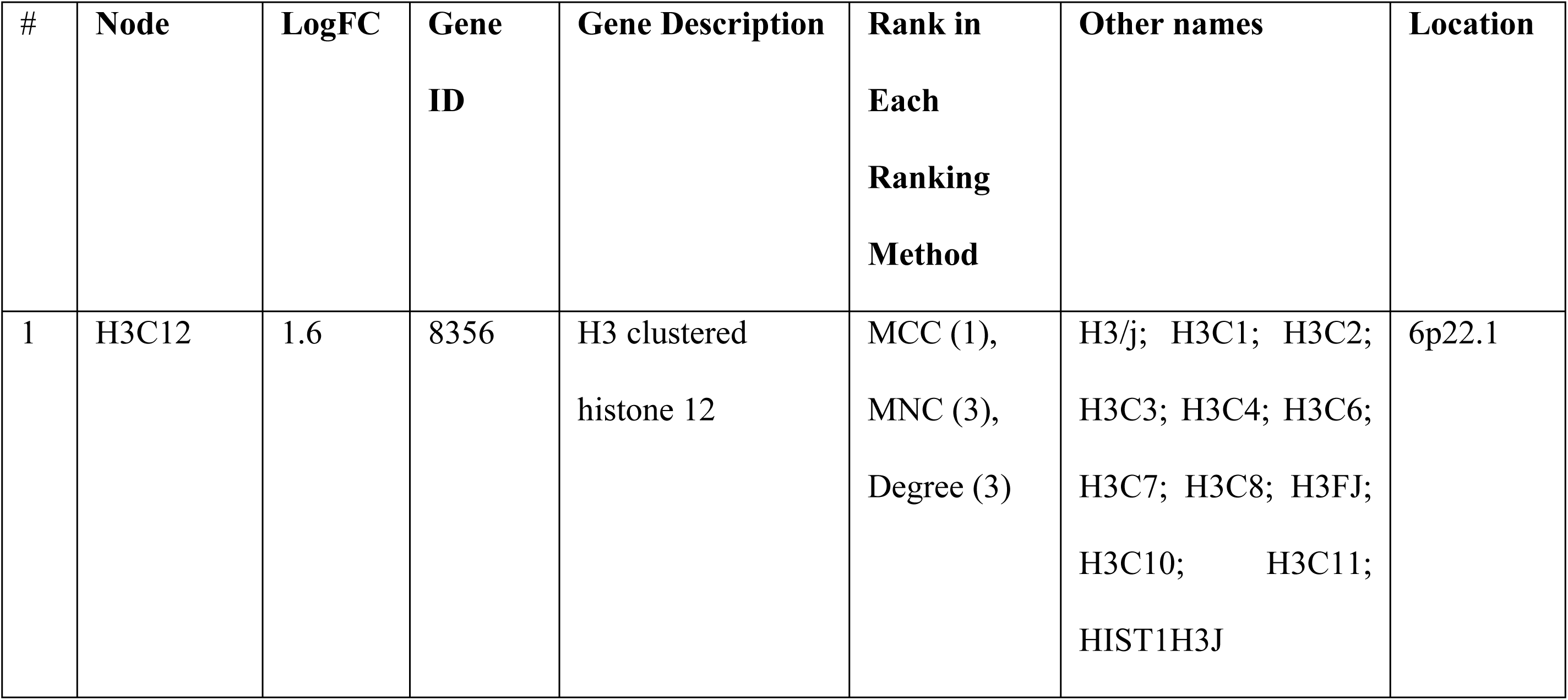
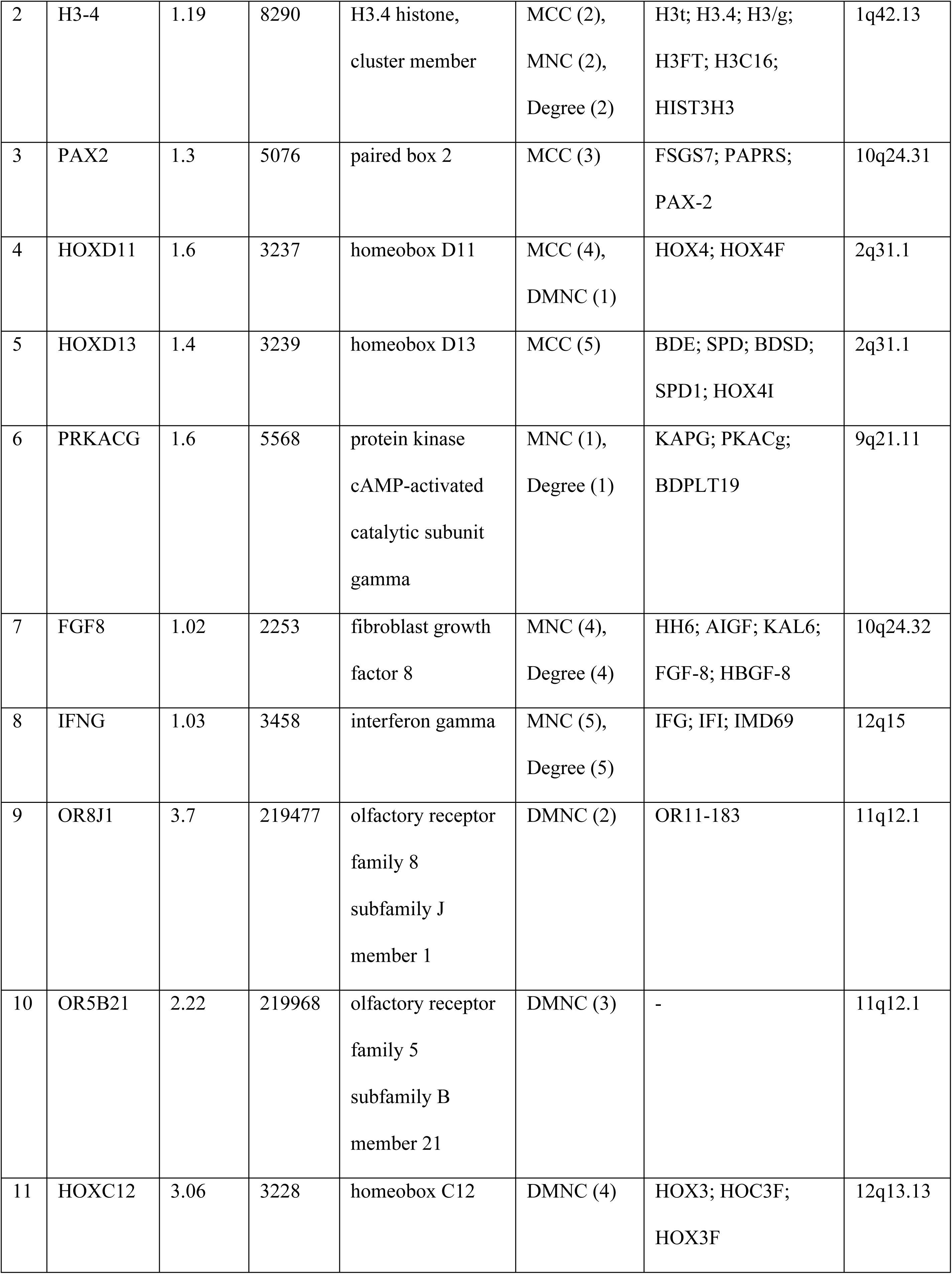
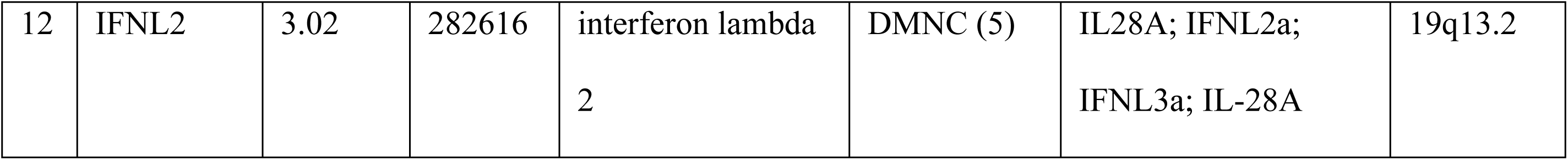
Hub genes from the PPI network of ∼6,500 DEGs, prioritized based on overlap across four cytoHubba topological methods. Gene locations and aliases are provided for annotation.

A second, focused network built from the top 200 DEGs (100 up, 100 down) yielded 93 nodes and 318 edges (**Fig 1**) and highlighted 13 hubs (HOXC8, HOXC9, HOXA11, HOXC11, HOXA13, WT1, ADIPOQ, GCG, VSTM2A, ATP4A, KCNE2, PTF1A) with both strong expression shifts and network centrality, supporting their potential as diagnostic/prognostic biomarkers (**Table 2**). This dual-network strategy separates broadly actionable targets from clinically tractable biomarkers by integrating systems-level topology with differential expression magnitude.

**Fig 1.**
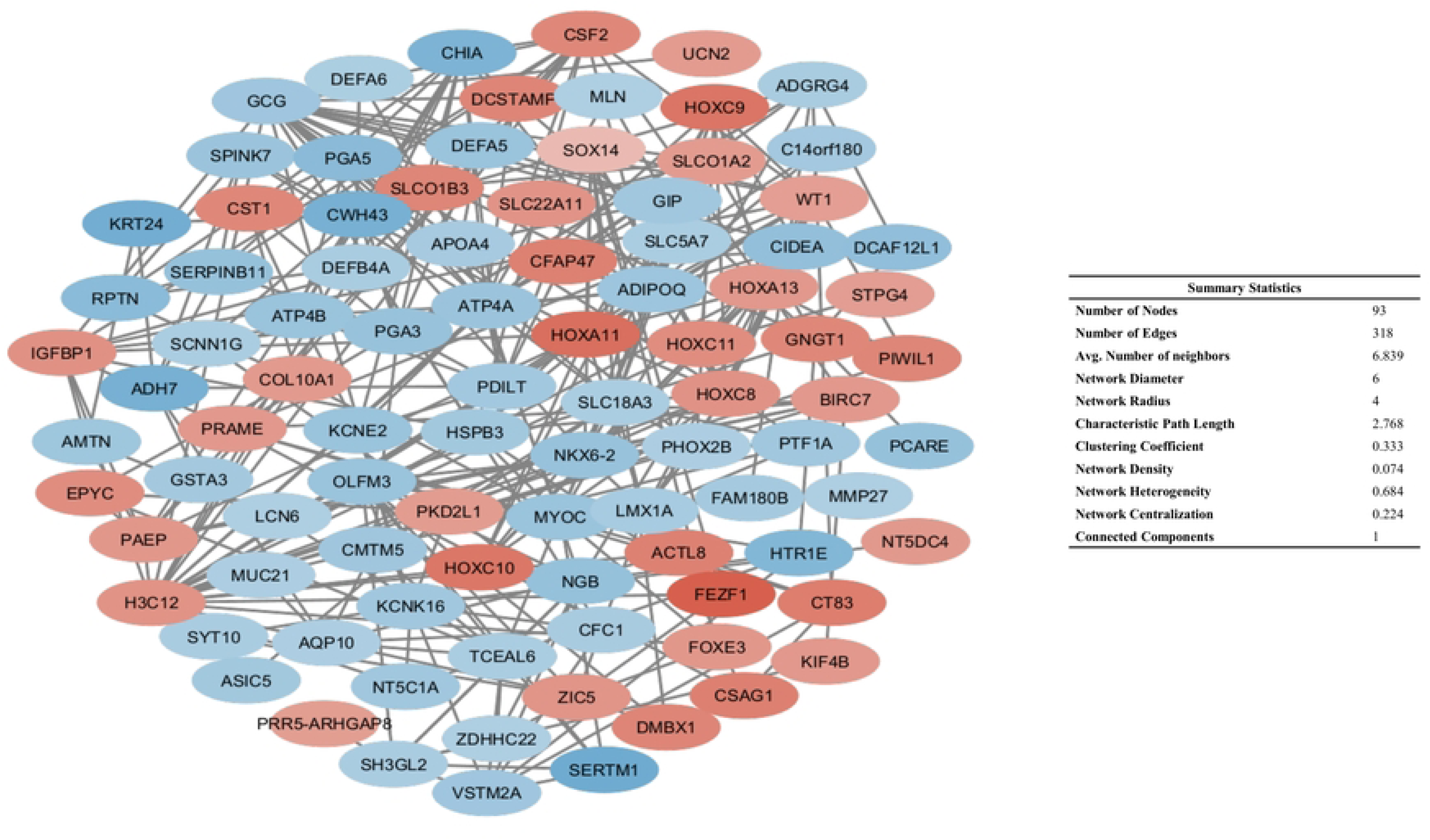
PPI network of the top 200 DEGs, with nodes colored by expression (red: upregulated, blue: downregulated). Constructed using STRING in Cytoscape (confidence score: 0.15).

**Table 2.** Hub genes from the PPI network of the top 200 DEGs, including ranking methods and log2 fold change (logFC) values.

| # | Node | LogFC | Gene ID | Gene Description | Rank | Ranking Method | Other names | Location |
| --- | --- | --- | --- | --- | --- | --- | --- | --- |
| 1 | H3C12 | 1.6 | 8356 | H3 clustered histone 12 | 1, 1, 1 | MCC, MNC, Degree | H3/j; H3C1; H3C2; H3C3; H3C4; H3C6; H3C7; H3C8; H3FJ; H3C10; H3C11; HIST1H3J | 6p22.1 |
| 2 | HOXC8 | 1.6 | 3224 | homeobox C8 | 2 | MCC | HOX3; HOX3A | 12q13.13 |
| 3 | HOXC9 | 2.1 | 3225 | homeobox C9 | 3 | MCC | HOX3; HOX3B | 12q13.13 |
| 4 | HOXA11 | 2.2 | 3207 | homeobox A11 | 4 | MCC | HOX1; HOX1I; RUSAT1 | 7p15.2 |
| 5 | HOXC11 | 1.7 | 3227 | homeobox C11 | 5, 1 | MCC, DMNC | HOX3H | 12q13.13 |
| 6 | HOXA13 | 1.5 | 3209 | homeobox A13 | 2 | DMNC | HOX1; HOX1J | 7p15.2 |
| 7 | PTF1A | -1.2 | 256297 | pancreas associated<br>transcription factor<br>1a | 3 | DMNC | p48; PACA;<br>PAGEN2;<br>bHLHa29; PTF1-p48 | 10p12.2 |
| 8 | VSTM2A | -1.2 | 222008 | V-set and<br>transmembrane<br>domain containing<br>2A | 4 | DMNC | VSTM2 | 7p11.2 |
| 9 | KCNE2 | -1.2 | 9992 | potassium voltage-<br>gated channel<br>subfamily E<br>regulatory subunit<br>2 | 5 | DMNC | LQT5; LQT6;<br>ATFB4; MIRP1 | 21q22.11 |
| 10 | GCG | -1.2 | 2641 | glucagon | 2, 2 | MNC,<br>Degree | GLP1; GLP2;<br>GRPP; GLP-1 | 2q24.2 |
| 11 | WT1 | 1.4 | 7490 | WT1 transcription<br>factor | 3, 4 | MNC,<br>Degree | GUD; AWT1;<br>WAGR; WT-1;<br>WT33; NPHS4;<br>WIT-2 | 11p13 |
| 12 | ADIPOQ | -1.2 | 9370 | adiponectin, C1Q<br>and collagen<br>domain containing | 3, 5 | MNC,<br>Degree | ACDC; ADPN;<br>APM1; APM-1;<br>GBP28; ACRP30;<br>ADIPQTL1 | 3q27.3 |
| 13 | ATP4A | -1.2 | 495 | ATPase H<sup>+</sup>/K<sup>+</sup><br>transporting subunit<br>alpha | 3, 3 | MNC,<br>Degree | ATP6A | 19q13.12 |

### 3.3. Functional enrichment analysis

Functional enrichment analysis was performed to understand the biological role of identified up-and downregulated genes. GO terms and pathway associations were analyzed in two sets separately to disclose different functional profiles of both gene sets (**Fig 2** and **3**).

**Fig 2.**
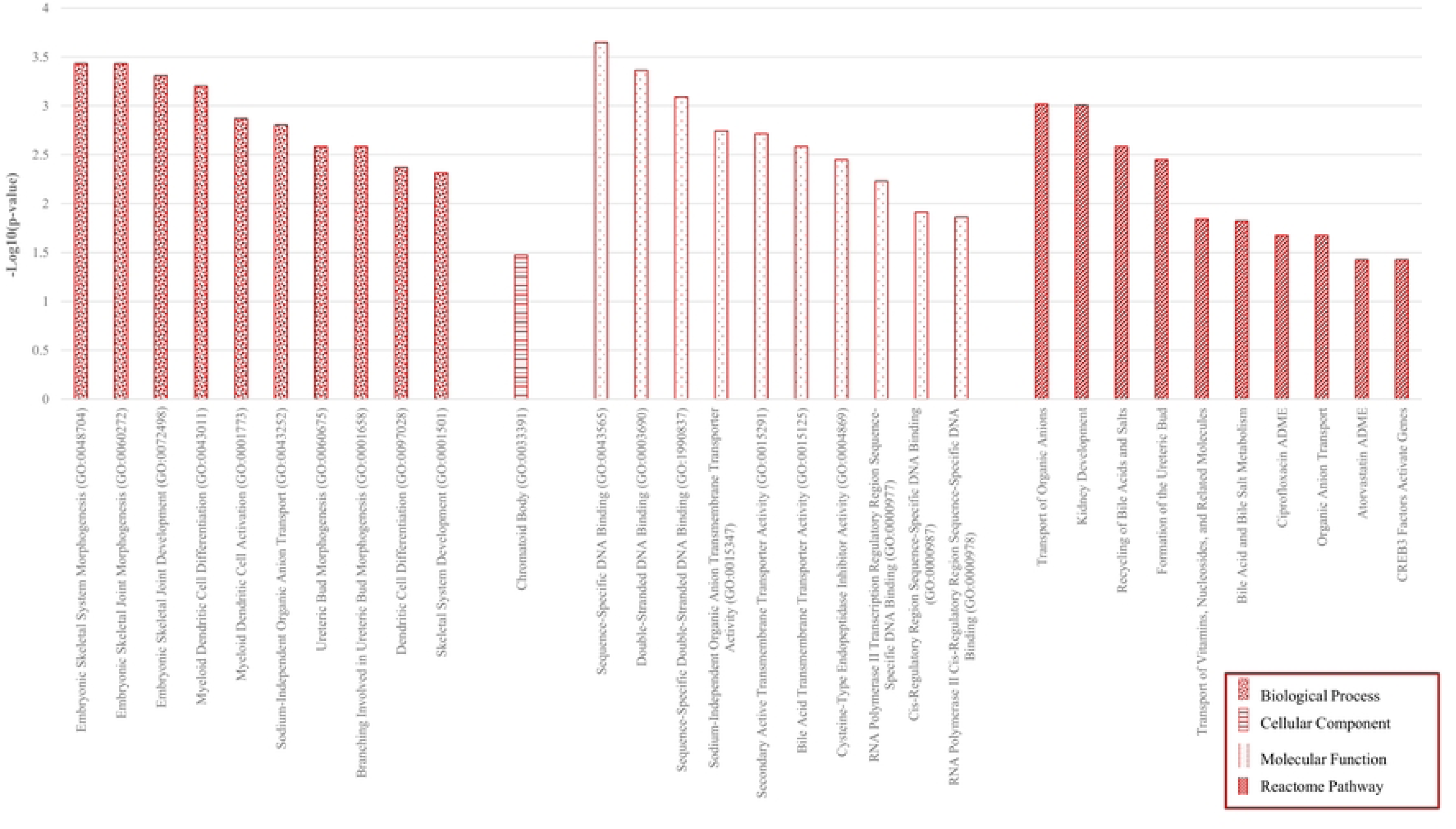
Functional enrichment analysis of upregulated genes. Top GO terms (BP, CC, MF) and Reactome pathways enriched among 85 upregulated GC genes, ranked by -Log_10_FDR using ENRICHR.

**Fig 3.**
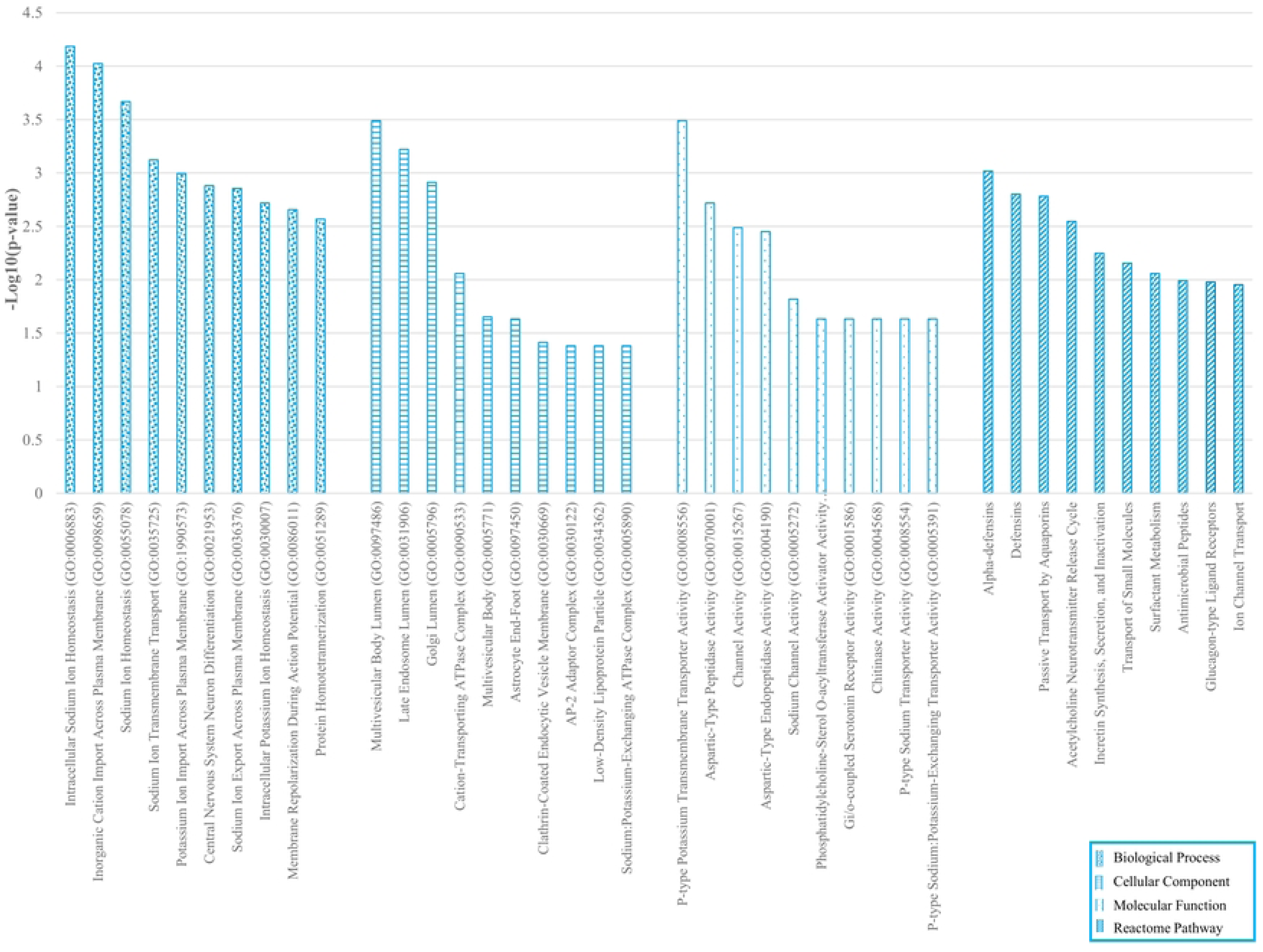
Functional enrichment of downregulated genes. Top GO terms and Reactome pathways for 94 downregulated GC genes, ranked by -Log_10_FDR using ENRICHR.

Upregulated genes (n = 85) were enriched for embryonic/developmental programs and immune differentiation (e.g., morphogenesis terms and myeloid dendritic cell activation), along with transcriptional regulation (sequence-specific DNA binding) and multiple solute/anion transport activities. Reactome analysis similarly highlighted organic anion and vitamin/nucleoside transport, kidney developmental pathways, and drug-metabolism modules (e.g., ciprofloxacin/atorvastatin ADME, bile acid metabolism), supporting a phenotype of developmental reactivation, transcriptional control, and altered transport/metabolic capacity.

Downregulated genes were predominantly enriched for ion homeostasis and membrane transport, particularly sodium/potassium handling (transmembrane transport, export/import, membrane repolarization), and localized to vesicular/endosomal–Golgi compartments and transport complexes (e.g., clathrin-coated vesicles, Na⁺/K⁺-ATPase complexes). Functional terms also indicated reduced channel and transporter activities, with Reactome showing decreased innate immune defense (defensins/antimicrobial peptides), aquaporin and ion channel transport, and neuroendocrine signaling pathways (acetylcholine release, incretin/glucagon signaling), consistent with loss of normal physiological and immune functions.

Overall, GC displayed a shift toward developmental reprogramming, transcriptional activation, and altered solute/drug metabolism (upregulated genes), alongside suppression of epithelial ion/transport homeostasis and innate immunity (downregulated genes), underscoring tissue remodeling and functional derangement relevant to biomarker and target discovery.

### 3.4. Cluster analysis and reactome pathway enrichment

We further enriched the top four clusters obtained from IPCA into Reactome pathways to gain deeper insights into the modular organization of the gene network and to identify distinct biological themes. These clusters were mainly enriched for FGFR signaling and its canonical downstream pathways, developmental biology, and transcriptional regulation, showing that they are biologically coherent and functionally specialized (**Fig 4**, and **Table 3**).

**Fig 4.**
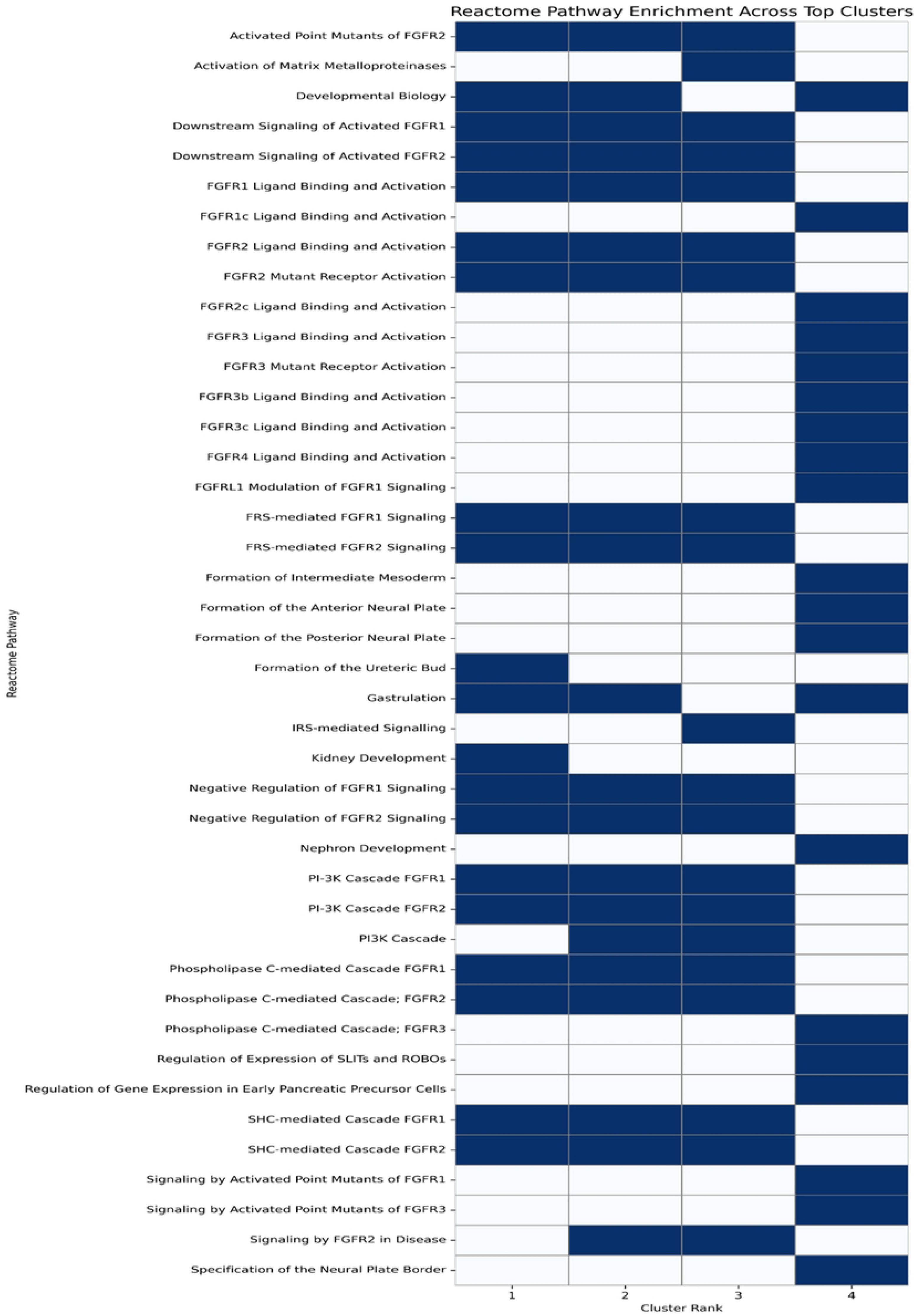
Heatmap of Reactome pathway enrichment across top-ranked clusters. The presence of a pathway in a given cluster is indicated in blue, and absence is shown in white. Data are based on the top 20 enriched Reactome pathways for each of the top four PPI network clusters identified using the IPCA algorithm in CytoCluster.

**Table 3.** Classification of Reactome Pathways Enriched Across Top Four Clusters.

| Category | Reactome Pathway | Clusters |
| --- | --- | --- |
| FGFR Signaling | FGFR1 Ligand Binding and Activation | 1, 2, 3 |
|  | FGFR1c Ligand Binding and Activation | 4 |
|  | FGFR2 Ligand Binding and Activation | 1, 2, 3 |
|  | FGFR2c Ligand Binding and Activation | 4 |
|  | FGFR3 Ligand Binding and Activation | 4 |
|  | FGFR3b Ligand Binding and Activation | 4 |
|  | FGFR3c Ligand Binding and Activation | 4 |
|  | FGFR4 Ligand Binding and Activation | 4 |
|  | FGFR2 Mutant Receptor Activation | 1, 2, 3 |
|  | FGFR3 Mutant Receptor Activation | 4 |
|  | Activated Point Mutants of FGFR2 | 1, 2, 3 |
|  | Activated Point Mutants of FGFR1 | 4 |
|  | Activated Point Mutants of FGFR3 | 4 |
|  | Signaling by FGFR2 in Disease | 2, 3 |
|  | FGFRL1 Modulation of FGFR1 Signaling | 4 |
|  | Negative Regulation of FGFR1 Signaling | 1, 2, 3 |
|  | Negative Regulation of FGFR2 Signaling | 1, 2, 3 |
| Canonical Signaling<br>Cascades | SHC-mediated Cascade FGFR1 | 1, 2, 3 |
|  | SHC-mediated Cascade FGFR2 | 1, 2, 3 |
|  | FRS-mediated FGFR1 Signaling | 1, 2, 3 |
|  | FRS-mediated FGFR2 Signaling | 1, 2, 3 |
|  | Phospholipase C-mediated Cascade FGFR1 | 1, 2, 3 |
|  | Phospholipase C-mediated Cascade; FGFR2 | 1, 2, 3 |
|  | Phospholipase C-mediated Cascade; FGFR3 | 4 |
|  | PI-3K Cascade FGFR1 | 1, 2, 3 |
|  | PI-3K Cascade FGFR2 | 1, 2, 3 |
|  | PI3K Cascade | 2, 3 |
|  | IRS-mediated Signalling | 3 |
|  | Downstream Signaling of Activated FGFR1 | 1, 2, 3 |
|  | Downstream Signaling of Activated FGFR2 | 1, 2, 3 |
| Developmental and<br>Organogenesis | Developmental Biology | 1, 2, 4 |
|  | Gastrulation | 1, 2, 4 |
|  | Kidney Development | 1 |
|  | Nephron Development | 4 |
|  | Formation of the Ureteric Bud | 1 |
|  | Formation of Intermediate Mesoderm | 4 |
|  | Formation of the Anterior Neural Plate | 4 |
|  | Formation of the Posterior Neural Plate | 4 |
|  | Specification of the Neural Plate Border | 4 |
| Transcriptional Regulation | Regulation of Gene Expression in Early Pancreatic Precursor Cells | 4 |
|  | Regulation of Expression of SLITs and ROBOs | 4 |
| Matrix Remodeling and Invasion | Activation of Matrix Metalloproteinases | 3 |

Across all clusters, FGFR signaling emerged as a consistent hallmark, including ligand-driven activation of FGFR1–4 and isoforms (FGFR1c/2c/3b/3c), alongside evidence of oncogenic FGFR2 mutant activation and regulatory counterbalance via negative modulators (e.g., FGFRL1). Enrichment extended to major downstream cascades, SHC/FRS-mediated signaling, PLC, and PI3K/IRS pathways, implicating proliferation, survival, motility, and metabolic regulation. Developmental programs were also reactivated (e.g., gastrulation, kidney/nephron and ureteric bud development, neural plate formation), suggesting increased cellular plasticity and possible EMT-related microenvironmental shifts. Additional cluster-specific themes included disrupted transcriptional control (early pancreatic precursor programs; SLIT/ROBO regulation) and matrix remodeling via metalloproteinase activation. Overall, the IPCA clusters define a coherent network in which FGFR signaling links developmental reprogramming, intracellular signaling, transcriptional regulation, and invasive behavior in gastric cancer.

### 3.5. Analysis of tumor grade-specific gene expression and potential biomarker utility

Gene expression analysis across tumor grades in stomach adenocarcinoma (STAD) uncovered a dynamic transcriptional landscape, with several genes exhibiting significant and grade-specific expression changes (**Fig 5**). These alterations offer insight into molecular mechanisms underlying tumor progression and differentiation, and highlight potential diagnostic and prognostic biomarkers.

**Fig 5.**
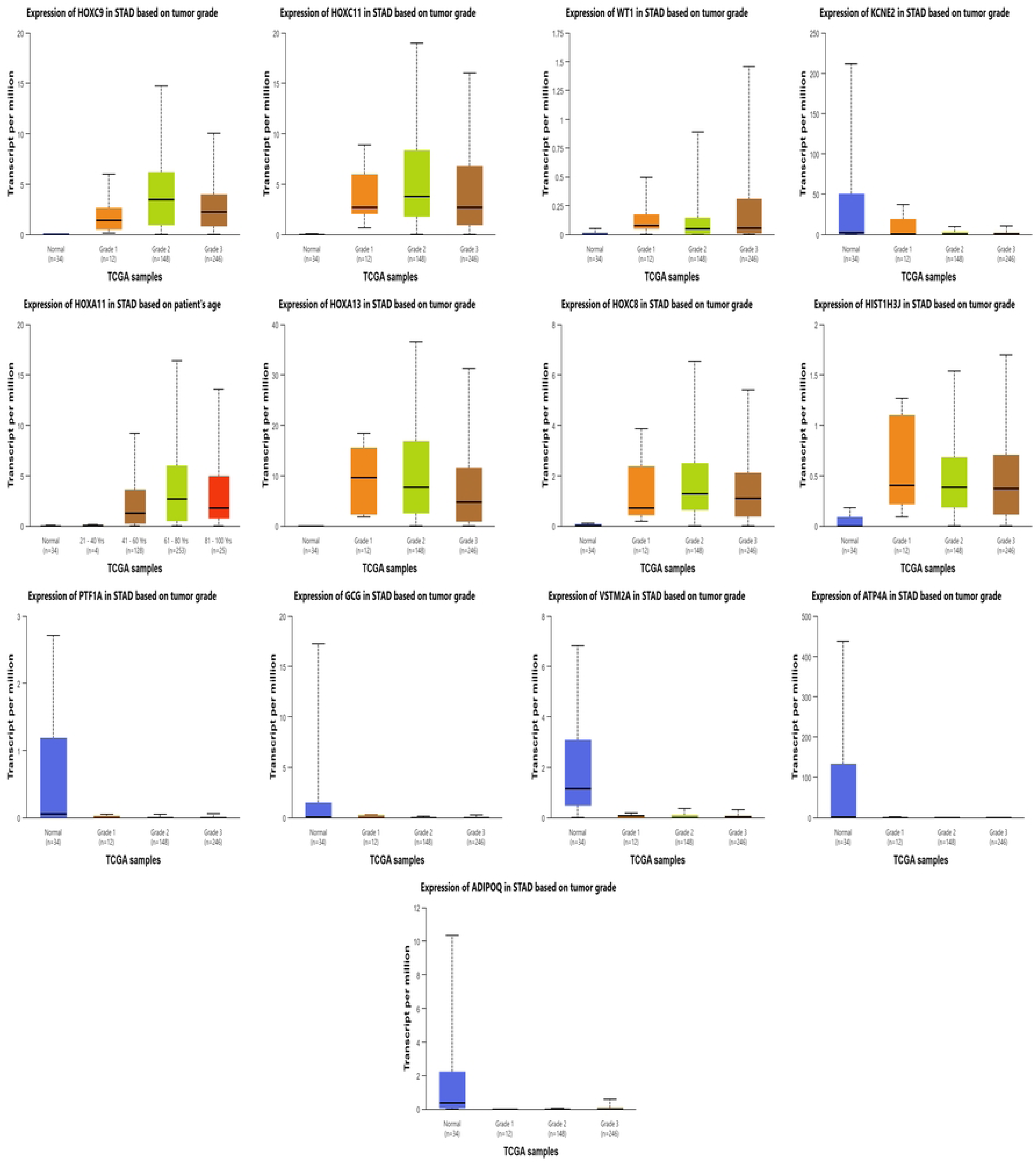
Grade-specific expression of 13 candidate genes in GC. Boxplots show transcript per million (TPM) values across normal and cancer grades 1-3 (TCGA data via UALCAN).

Grade-stratified analysis showed increasing expression of HOXA11/HOXA13/HOXC8/HOXC9/HOXC11 and HIST1H3J from Grade 1 to Grades 2–3, consistent with developmental/epigenetic reprogramming. In contrast, gastric lineage markers (ATP4A, KCNE2, PTF1A, VSTM2A) were strongly downregulated early and remained low across grades, indicating sustained loss of epithelial identity. WT1 rose mainly in Grades 2–3, while ADIPOQ was repressed early with modest recovery and GCG declined gradually. Together, these patterns support a panel where HOX/HIST1H3J (±WT1) reflect progression, and ATP4A/KCNE2/PTF1A/VSTM2A (±ADIPOQ) mark early dedifferentiation.

### 3.6. Expression of genes in STAD based on nodal metastasis status and biomarker potential

When TCGA-STAD samples were examined according to nodal metastasis stage (N0–N3), a clear and recurring alteration appeared among the 13 studied genes, pointing to their possible value as biomarkers for identifying and tracking disease (**Fig 6**).

**Fig 6.**
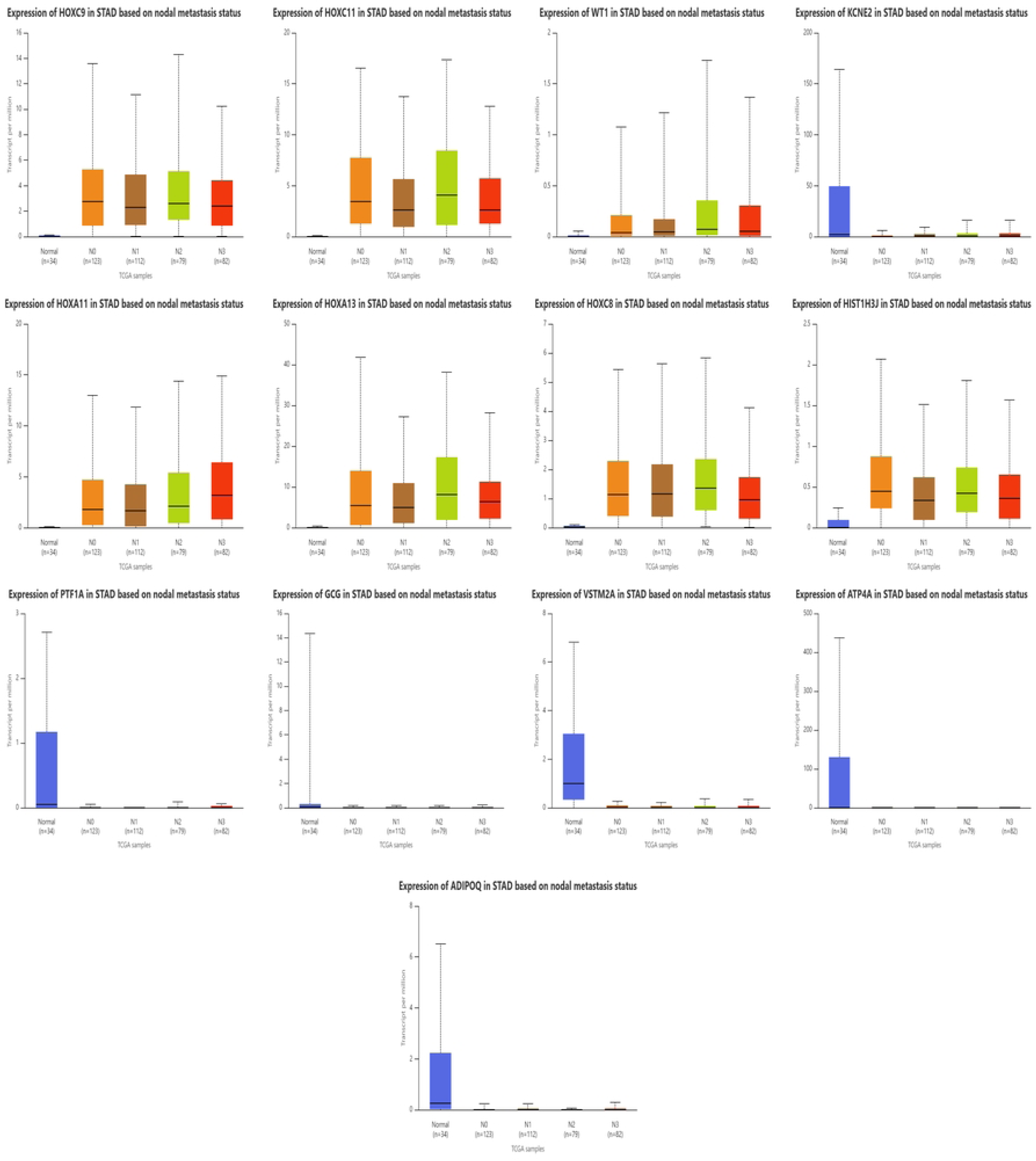
Transcript expression levels of selected genes in GC patients based on nodal metastasis status.

Early Detection and Diagnostic Biomarkers: The HOX cluster genes (HOXA11, HOXA13, HOXC8, HOXC9, and HOXC11) together with HIST1H3J, showed marked and statistically strong overexpression across every nodal category when compared with normal gastric tissue (most with P ≤ 10⁻⁹). The elevated expression was already present in node-negative tumors, pointing to early engagement of these transcription factors in tumor development rather than secondary activation during metastasis. Seen in this light, their persistent activity may serve as an early warning sign for primary tumors that have yet to reach the lymphatic system.

Progression and Metastatic Biomarkers: While the expression of HOX genes and HIST1H3J did not show a progressive increase with higher nodal stages (N1–N3), certain genes demonstrated patterns suggestive of progression markers. Expression of WT1 rose sharply in N0 and N1 tumors (P around 10⁻⁶–10⁻⁷) and climbed even higher by the N3 stage, which may point to its role in promoting or tracking advanced nodal spread. The smaller but steady changes seen for KCNE2 and PTF1A across the same stages might reflect the gradual erosion of differentiation as tumors become more metastatic, an observation that deserves closer study for its potential prognostic value.

Loss of Differentiation Markers: Gastric lineage genes (ATP4A, KCNE2, PTF1A, VSTM2A) and ADIPOQ were markedly downregulated in both node-negative and metastatic tumors (all P < 0.01 vs normal), indicating early, sustained dedifferentiation and supporting their use as negative diagnostic markers independent of nodal status. GCG showed no nodal-group differences and can be excluded from the core biomarker set. Overall, the data support a two-tier framework: early activation markers (HOX-cluster genes, HIST1H3J), persistent loss-of-identity markers (ATP4A, KCNE2, PTF1A, VSTM2A, ADIPOQ), and WT1 as a potential progression marker associated with nodal spread.

### 3.7. Tumor stage–specific gene expression and implications for biomarker discovery in STAD

As we looked across tumor stages in STAD, certain genes stood out for how sharply their expression shifted; patterns that could eventually guide biomarker discovery and treatment design. Several genes, among them HOXA11, HOXA13, HOXC8, HOXC9, HOXC11, HIST1H3J, WT1, and the differentiation markers ATP4A, KCNE2, VSTM2A, and PTF1A, showed stage-specific shifts in activity, suggesting value for classifying tumors or refining clinical decisions (**Fig 7**).

**Fig 7.**
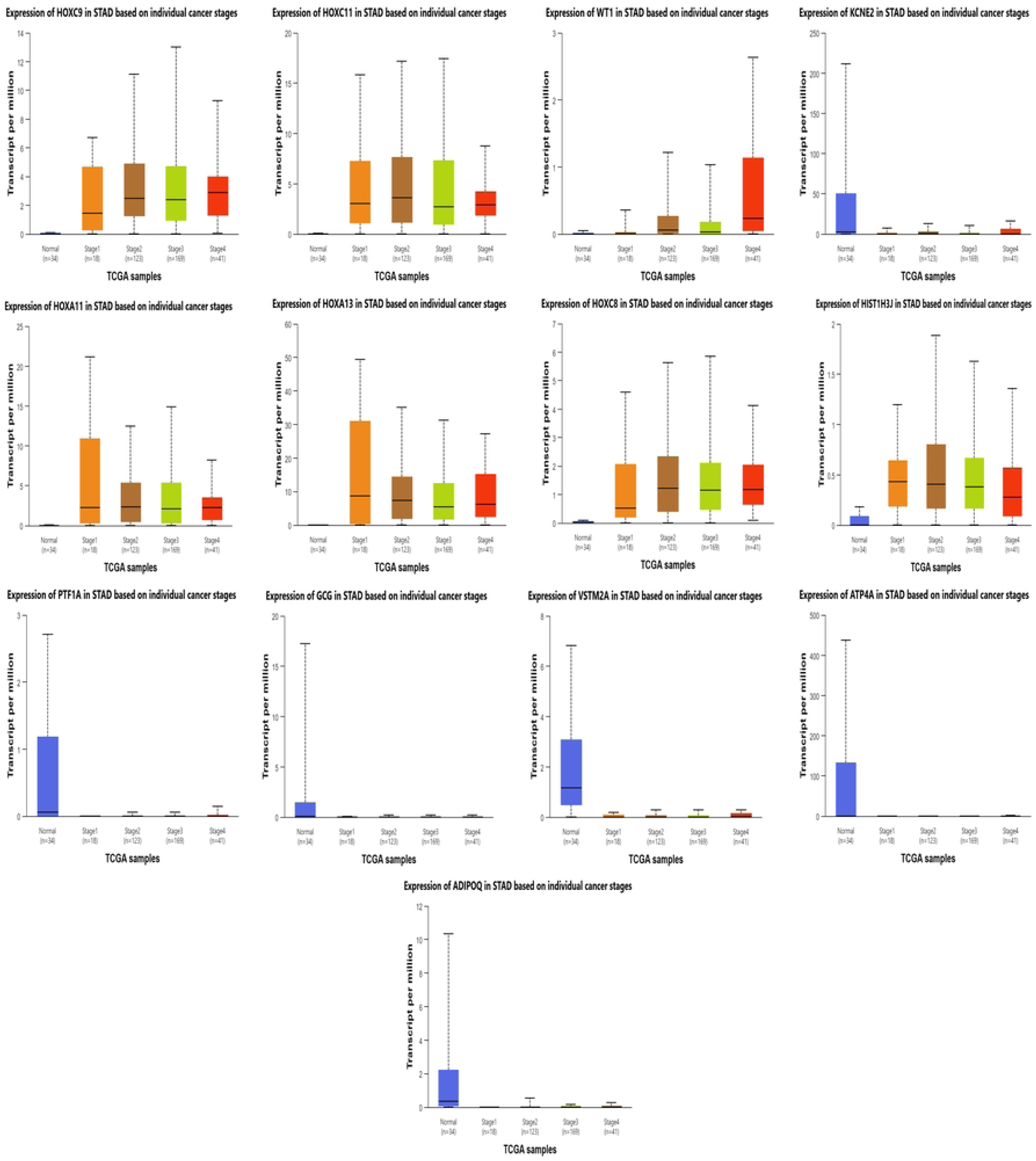
Stage-specific expression of 13 candidate genes in GC. Boxplots show transcript per million (TPM) values across normal tissue and cancer stages 1-4 (TCGA data via UALCAN).

Across STAD stages, HIST1H3J and multiple HOX-cluster genes are consistently upregulated (often P < 10⁻¹²), with early onset and sustained expression, suggesting roles in tumor establishment and maintenance and potential utility as early molecular markers. In contrast, gastric differentiation genes ATP4A, KCNE2, VSTM2A, and PTF1A are persistently downregulated across all stages versus normal tissue, indicating stable loss of parietal/ductal identity and supporting their use as diagnostic markers of malignant transformation. WT1 increases with stage, reaching stronger significance in advanced disease, consistent with a progression-associated biomarker. Together, these patterns define a stable STAD signature, early HOX/HIST1H3J activation with sustained repression of gastric identity genes, while WT1 may aid risk stratification and monitoring.

### 3.8. Survival analysis of candidate genes in STAD

We used Kaplan–Meier analysis to examine how the 13 candidate genes relate to patient survival in STAD. Of all the genes tested, only ADIPOQ showed a significant link with overall survival (p = 0.012); the others showed no clear association (p > 0.05) (**Fig 8**).

**Fig 8.**
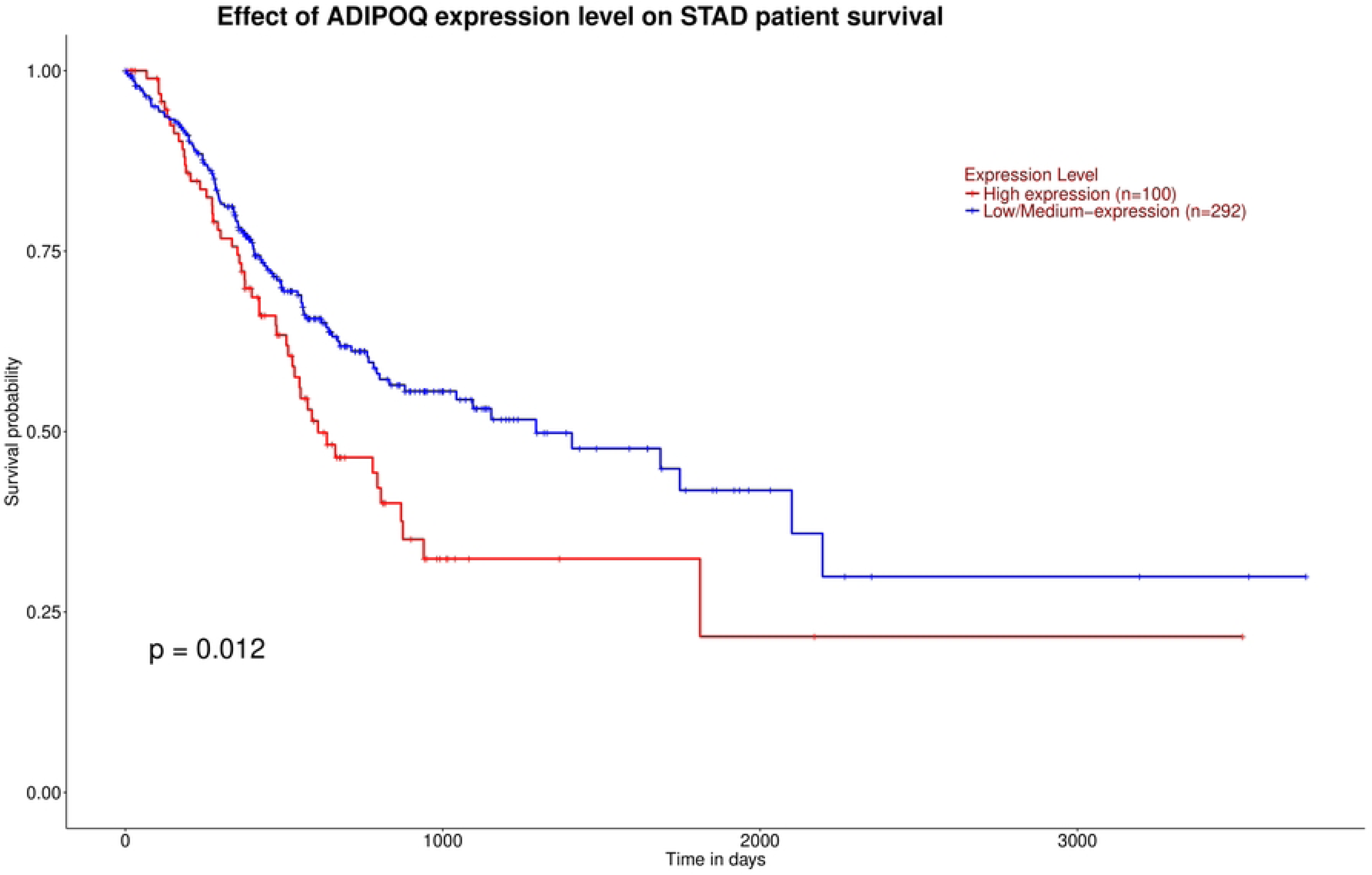
Survival analysis of ADIPOQ in GC patients.

Patients were stratified by ADIPOQ expression (high, n = 100 vs low/intermediate, n = 292). Kaplan–Meier analysis showed poorer survival in the high-expression group, suggesting ADIPOQ as a negative prognostic marker in STAD. As adiponectin modulates metabolism and inflammation, this pattern supports a context-dependent role in the tumor microenvironment. Clinically, ADIPOQ may aid risk stratification, but independent cohort validation and mechanistic studies are needed.

## 4. Discussion

In this work, we took a broad look at the transcriptome of STAD to map gene expression changes and their biological and clinical implications. By combining a full set of roughly 6,500 DEGs with a focused analysis of the top 200, two clear interaction modules began to emerge from the data. The first includes histone genes, immune regulators, and kinases that occupy central positions in the protein–protein interaction network, making them attractive targets for therapeutic disruption. The second group includes transcription factors, epithelial genes, and metabolic regulators that show striking expression differences. These features make them appealing candidates for developing both diagnostic and prognostic markers.

We found that HOXA11, HOXA13, HOXC8, HOXC9, and HOXC11 were consistently more active across tumor grades, stages, and nodal groups, a result that fits well with earlier evidence of HOX gene reactivation in gastrointestinal cancers [20]. Their expression tended to climb as tumors became less differentiated and more advanced, most noticeably in the aggressive forms. This steady rise links them to both loss of cellular identity and increasing malignancy. The histone gene HIST1H3J showed a comparable pattern, its levels rose step by step with tumor grade and stage, which aligns with reports of broad epigenetic remodeling [21]. These genes seem to switch on early in tumor formation and remain active as the disease develops, helping to sustain transcriptional activity that supports growth and survival.

Conversely, the observed downregulation of gastric lineage markers (ATP4A, KCNE2, VSTM2A, PTF1A) from early stages supports the idea that dedifferentiation is an initiating event in gastric tumorigenesis [22, 23]. Our observations agree with earlier reports showing that genes tied to acid secretion and epithelial differentiation are already dampened in the early stages of gastric cancer. In particular, several studies have noted reduced expression of ATP4A and ATP4B, both of which have been examined as potential diagnostic markers [22]. ADIPOQ also showed dynamic regulation, with an early decrease followed by partial reactivation in advanced tumors; recent work suggests that low adiponectin levels or expression correlate with more aggressive GC features [24]. Survival analysis revealed that high ADIPOQ expression correlates with poorer patient outcomes, further highlighting its potential as a context-dependent prognostic biomarker [24].

Our enrichment analysis pointed to a set of molecular patterns that mirror what other transcriptomic studies have described in gastric cancer. The genes showing higher expression were mostly involved in transcriptional control, immune signaling, and developmental processes, similar to earlier reports describing a reactivation of embryonic and inflammatory pathways during tumor growth [25]. By contrast, the genes showing reduced activity were mostly linked to ion transport, vesicle movement, and antimicrobial defense. Their decline mirrors the loss of normal gastric cell function that tends to accompany malignant transformation [26]. Overall, the data suggest a broad reorganization of the tumor transcriptome, shifting away from the physiological roles of the stomach toward a more adaptable, development-like and metabolically flexible state, a trend that recent single-cell and spatial transcriptomic analyses of gastric and gastrointestinal cancers have also captured [25, 26]. Our cluster-based pathway enrichment further emphasized the centrality of FGFR signaling and its downstream cascades. This finding is in agreement with recent clinical and preclinical studies demonstrating FGFR dysregulation as a driver of oncogenic transcriptional programs in gastric cancer [27, 28].

A key strength of this study is its layered design, combining stringent DEG filtering with network modeling, functional annotation, and clinicopathological interpretation to identify clinically relevant targets. Our findings are consistent with recent bulk and single-cell GC studies reporting metabolic reprogramming, immune microenvironment changes, and loss of gastric differentiation [7, 29]. Limitations include reliance on TCGA-STAD only, lack of experimental validation, and the inability of transcriptomics to capture post-transcriptional/post-translational regulation; future work should validate results in independent cohorts and integrate additional omics layers.

Going forward, it will be important to check how well these biomarkers perform in other patient cohorts and to see whether they can be tracked in easier-to-obtain materials like blood or gastric fluid. Testing how HOX genes, WT1, and FGFR-pathway effectors behave when perturbed in cells or organoids should also help clarify what roles they actually play in tumor biology. Additionally, the relationship between ADIPOQ expression and the immune microenvironment warrants investigation to decipher its dual role in metabolism and immunoregulation.

In summary, our study delineates two functionally distinct gene modules in gastric cancer: a regulatory core enriched in HOX and histone genes that may be exploited for early diagnosis and a set of suppressed differentiation genes marking the loss of gastric identity. These results outline a molecular framework that could guide biomarker-based patient classification and help shape new therapeutic strategies in STAD. They also highlight how combining network-level analysis with clinical data can move the field closer to truly precise cancer care.

## 5. Conclusion

Using an integrated transcriptomic and network-based framework (differential expression, PPI mapping, hub ranking, and pathway enrichment), we identified a coordinated STAD signature marked by upregulation of developmental/immune programs and repression of epithelial differentiation and transport genes. Dual-network analysis prioritized a focused biomarker set—HOX genes, HIST1H3J, ATP4A, KCNE2, and PTF1A—with potential utility for early detection, monitoring, and risk stratification, and highlighted actionable pathways relevant to precision oncology. Key limitations are reliance on TCGA-STAD data and lack of experimental validation; future work should validate these markers in independent cohorts and test mechanisms in vitro/in vivo, ideally integrating proteomic/epigenomic and spatial transcriptomic layers to refine heterogeneity and clinical translation.

## Abbreviations

BP: Biological Process
CC: Cellular Component
DEG: Differentially Expressed Gene
FGFR: Fibroblast Growth Factor Receptor
GC: Gastric Cancer
GO: Gene Ontology
HOX: Homeobox Gene
KEGG: Kyoto Encyclopedia of Genes and Genomes
MF: Molecular Function
PPI: Protein–Protein Interaction
RNA-seq: RNA Sequencing
STAD: Stomach Adenocarcinoma
TCGA: The Cancer Genome Atlas;
UALCAN: University of Alabama at Birmingham Cancer Data Analysis Portal.

## Declarations

### Ethics statement

This study used only publicly available, de-identified data and did not require ethics approval.

### Consent for publication

Not applicable.

### Data Availability

TCGA-STAD RNA-seq and clinical data are publicly available from the Genomic Data Commons (GDC). All processed data supporting the findings of this study are available at GitHub (https://github.com/negmot/tcga-stad-network-analysis) and have been archived on Zenodo (DOI: 10.5281/zenodo.18537980).

### Author contributions

N.M.D.: Conceptualization, Supervision, Data curation, Formal analysis, Visualization, Writing-original draft.

M.S.R.R.: Data curation, Investigation, Writing-review & editing.

Both authors: Final approval of the manuscript.

## Supporting information

**Supplementary File 1**. Full differential expression results tables (RAR/XLSX).

